# The kinase Rio1 and a ribosome collision-dependent decay pathway survey the integrity of 18S rRNA cleavage

**DOI:** 10.1101/2022.07.29.501969

**Authors:** Melissa D. Parker, Adam J. Getzler, Katrin Karbstein

## Abstract

The 18S rRNA sequence is highly conserved, particularly at its 3’-end. In contrast, the sequence around the 3’-end is degenerate with similar sites nearby. How RNA is correctly processed by the endonuclease Nob1 is not known, especially because *in vitro* experiments have shown it to be error-prone. Here we used yeast genetics, biochemistry, and next generation sequencing to investigate a role for Rio1 in monitoring the 3’-end of 18S rRNA. We demonstrate that Nob1 can miscleave its rRNA substrate and that miscleaved rRNA accumulates upon bypassing the Rio1-mediated quality control step, but not in healthy cells with intact quality control mechanisms. Mechanistically, we show that Rio1 binding to miscleaved rRNA is weakened. Accordingly, excess Pno1 results in accumulation of miscleaved rRNA. Ribosomes containing miscleaved rRNA enter the polysomes and produce dominant negative growth defects, suggesting that they cause defects during translation. Our data strongly suggest that ribosome collisions identify these miscleaved 18S rRNA-containing ribosomes as partially functional and target them for degradation. Altogether, the data support a model in which Rio1 inspects the 3’-end of the nascent 18S rRNA, only removing Nob1 and Pno1 from the ribosomes with precisely cleaved 18S rRNA to prevent miscleaved 18S rRNA-containing ribosomes from erroneously engaging in translation, where they induce ribosome collisions. The data also demonstrate how ribosome collisions “purify” the cells of altered ribosomes with different functionalities, with important implications for the concept of ribosome heterogeneity.

## Introduction

Ribosomes are the molecular machines responsible for protein synthesis in all cells. Maintaining translation fidelity and ensuring protein homeostasis requires proper ribosome assembly, which involves the transcription of 4 ribosomal RNAs (rRNAs), coupled to rRNA processing, folding, and binding to 79 ribosomal proteins (RPs) in a series of ordered steps involving over 200 transiently binding assembly factors^1,2^. During the final cytoplasmic assembly steps of the small ribosomal subunit (40S), cells have established a series of quality control (QC) mechanisms regulated by assembly and translation factors to probe the structural integrity and function of nascent ribosomes^3–5^. These QC checkpoints are important for maintaining healthy cells, as numerous cancers accumulate mutations that bypass ribosome QC^6–8^ and have altered RP stoichiometry leading to ribosome heterogeneity^9–11^. In addition to reducing ribosome abundance, haploinsufficiency of RPs can result in misassembled ribosomes lacking these RPs and predisposes patients to cancer^11–16^.

In the final stages of 40S ribosome assembly in yeast, the 18S rRNA 3’-end is formed from the precursor 20S rRNA by the essential endonuclease Nob1^17–20^, promoted by its binding partner Pno1^21^. Immediately prior to Nob1-mediated 18S rRNA cleavage, the precursor 40S (pre-40S) subunit containing 20S rRNA is bound to two assembly factors, Nob1 and Pno1^22–27^, and lacks the RP Rps26 as its binding site overlaps that of Pno1 on the mature and nascent ribosomes, respectively^26,28,29^. Pno1 stabilizes Nob1 on the ribosome and Nob1 blocks mRNA recruitment, thus creating a QC checkpoint that blocks pre-40S from translation^7,21^. After Nob1-dependent 18S rRNA cleavage, the ATPase Rio1 removes both Nob1 and Pno1 from the nascent 40S subunit, allowing for the recruitment of mRNA and Rps26^7,26,27,30^. Therefore, Rio1 is responsible for monitoring whether 18S rRNA 3’-end cleavage has occurred, and only licensing ribosomes with mature 18S rRNA for translation^7^.

It is vital for cells to block these immature pre-40S ribosomes from participating in translation, as translating 20S pre-rRNA-containing ribosomes have reduced translational fidelity and do not support cell growth^7,18,28,31^. Interestingly, 18S rRNAs with as few as 3 nucleotides of precursor rRNA sequence retained at the 18S rRNA 3’-end do not support cell viability in yeast either^32^, suggesting not only that is cleavage important, but that it must be precise. However, Nob1 does not always identify the cleavage site correctly, as Nob1 frequently miscleaves its rRNA substrate *in vitro*^5,20,33,34^. How Nob1 recognizes its cleavage site remains unknown, as does whether Nob1 miscleaves endogenous 18S rRNA *in vivo*, whether cleavage accuracy affects ribosome function, and if so, whether cleavage accuracy is monitored to prevent miscleaved rRNA-containing ribosomes from translating.

In addition to QC during ribosome assembly, cells actively monitor translation, targeting aberrant mRNA, rRNA, and nascent peptides for degradation^35,36^. When a mutation in the 18S rRNA decoding site (18S:A1492C) renders the 40S ribosome unable to bind and decode incoming tRNA during translation^37,38^, this mutant 18S rRNA is degraded through the so-called 18S nonfunctional rRNA decay (18S NRD) pathway^39,40^. These stalled mutant ribosomes initiate a cascade of events initiated by collisions with the trailing ribosomes, which include mRNA and 18S rRNA decay, and which require an overlapping set of proteins (Asc1, Slh1, Hel2, Rps3 ubiquitination, Dom34, Xrn1, and the Ski complex)^36,40–43^. As 18S NRD has only been studied in the context of the decoding deficient 18S:A1492C mutant, which is not expected to enter the open reading frame (ORF), it is unknown whether 18S NRD targets all nonfunctional 18S rRNAs even those translating more slowly inside the ORF, and if the same factors are involved in all cases.

Using next generation sequencing, yeast genetics, and biochemical techniques, we show that miscleaved 18S rRNAs are formed *in vivo*, but that their fate is regulated by the Rio1-mediated QC step and a collision-dependent NRD-like pathway. Next generation sequencing of the 3’-ends of 18S rRNAs from wild type yeast cells show that about 2% of 18S rRNAs are miscleaved. Biochemical and genetic data demonstrate that truncated, miscleaved 18S rRNAs, even at these low concentrations, disrupt cell growth, because they engage in translation where their compromised activity leads to ribosome collisions with correctly matured rRNAs. The resulting complexes are targeted for ribosome decay involving the proteins Asc1, Hel2, Dom34 and Xrn1. We show that miscleaved 18S rRNAs are stabilized upon bypassing the Rio1-mediated QC step, implicating Rio1 in QC of correct Nob1 cleavage. Confirming this, ribosomes containing truncated, miscleaved 18S rRNAs retain Pno1, indicating that they cannot pass the Rio1-mediated checkpoint. Finally, Rio1 has a stronger binding affinity for 18S rRNAs with correct 3’-ends than miscleaved 3’-ends. Altogether, these data support a model in which Rio1 inspects the 3’-end of 18S rRNA, ensuring only ribosomes with accurately cleaved 18S rRNA are released into the translating pool. On the other hand, Rio1 will not remove Nob1 or Pno1 from ribosomes containing miscleaved 18S rRNA, thus restricting their translation. The data also demonstrate how dysfunctional ribosomes produced via leaky checkpoints are removed from the translating pool via NRD, thereby purifying cells of dysfunctional heterogenous ribosomes.

## Results

### Nob1 miscleaves pre-18S rRNA

The 3’-end of 18S rRNA is highly conserved and identical in organisms ranging from yeast to humans (Figure S1A), indicating the importance of this rRNA segment. Furthermore, maintaining faithful 18S rRNA cleavage during ribosome assembly is critical, as 18S rRNAs retaining as few as 3 additional nucleotides (nts) at its 3’-end do not support viability S. *cerevisiae*^32^. While important, accurate cleavage of the 3’-end of 18S rRNA during pre-rRNA processing may not be a simple task. Nob1 cleaves the premature (pre-) rRNA between two adenosines, and there are multiple pairs of adenosines nearby (Figure S1B) that Nob1 could potentially recognize and cleave. Consistently, *in vitro* experiments have shown that Nob1 frequently miscleaves its 18S rRNA substrate, resulting in multiple cleavage products^5,20,33,34^. What remains unknown is whether Nob1 (alone or in combination with other assembly factors) can identify and accurately cleave endogenous 18S rRNA *in vivo*.

To examine this, we performed 3’-RACE (rapid amplification of cDNA ends) sequencing on 18S rRNA to survey the 3’-ends of 18S rRNA. We grew yeast to early stationary phase (between OD_600_ 1.2 – 1.8), purified the 40S ribosomal subunits, and extracted the 18S rRNA. Next, a linker was ligated to the 3’-ends of the RNA to protect it from degradation and accurately identify the 3’-ends. This linker was used to prime reverse transcription, creating cDNAs that were subsequently converted into sequencing libraries using 18S rRNA-specific primers for analysis of 18S rRNA 3’-ends. Due to concerns that reverse transcription through the Dim1-dimethylation site in 18S rRNA (m^6^_2_A1781 and m^6^_2_A1782) would pose complications, we used a yeast strain containing a mutation in Dim1 (Dim1-E85A) that does not dimethylate 18S rRNA^44^, thus allowing us to sequence the final 40-60 nucleotides of 18S rRNA. We obtained 2.8-4 million reads per sample, with 96-99% of reads aligning to the 3’-end of 18S rRNA. Mapping these reads showed that ∼98% of 18S rRNAs terminate at the mature 3’-end (Figure S1B). Only a small percentage (∼2%) of RNAs are miscleaved, producing mostly slightly shortened and some lengthened 18S rRNA products. The most frequent (0.03%) miscleavage downstream of the canonical 3’-end occurred between two adenosines one nucleotide into ITS1, the precursor rRNA sequence 3’ to 18S rRNA (18S+1 nt), while the most abundant (0.8%) upstream miscleavage occurred between a cytidine and an adenosine four nucleotides upstream of site D (18S-4 nts, Figure 1A-B). Thus, these data indicate a propensity for Nob1 to cleave 5’ to an adenosine but show that most 18S rRNAs have a correctly formed 3’-end.

**Figure 1:**
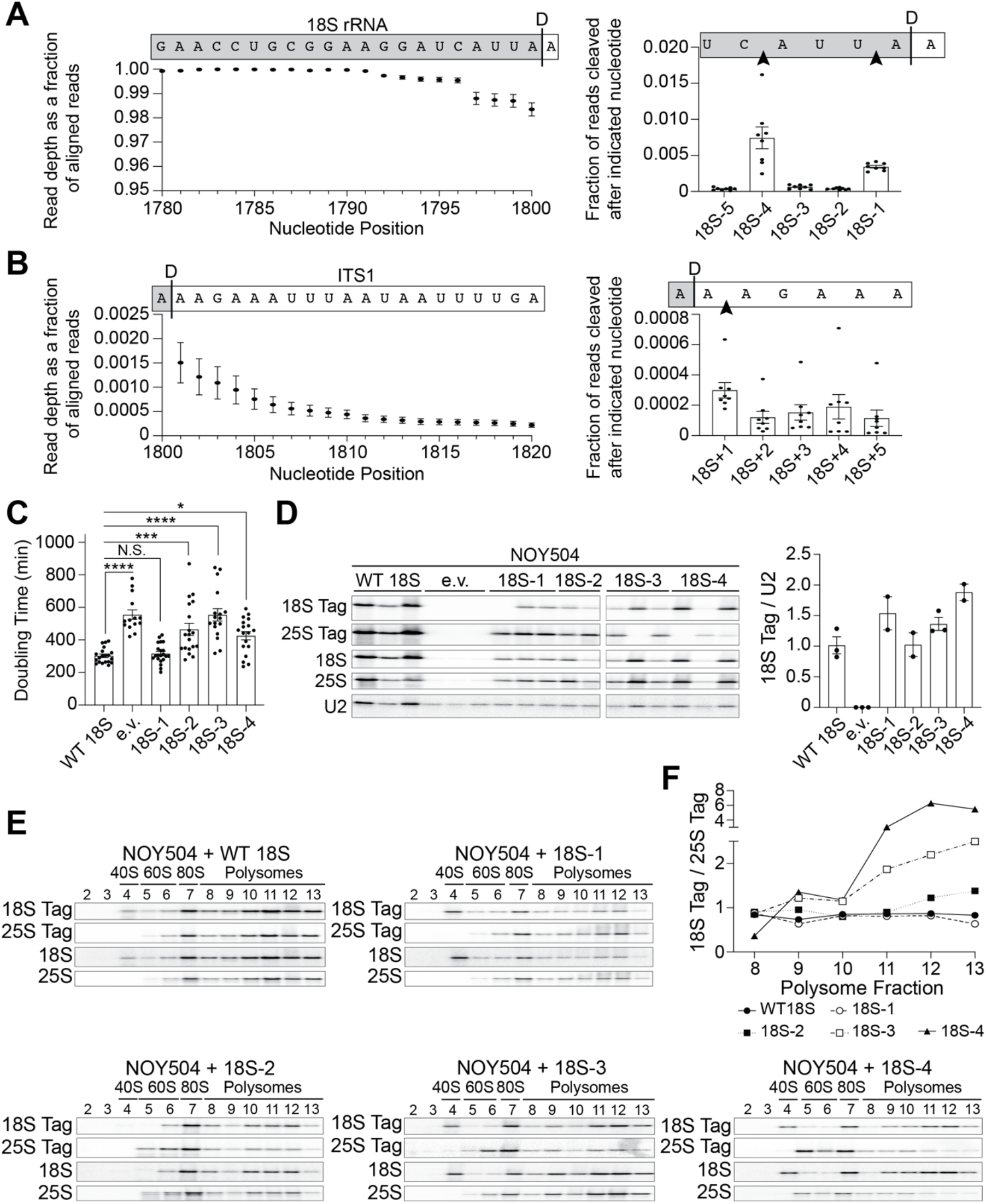
Miscleaved 18S rRNA are low in abundance but detrimental to cell growth. (A-B) 3’-RACE-sequencing of 18S rRNA extracted from 40S subunits purified from Gal::Pno1;Gal::Dim1 cells grown in glucose to deplete endogenous Pno1 and Dim1 and supplemented with plasmids expressing Pno1 and Dim1-E85A, an inactive mutant that prevents dimethylation of 18S rRNA^44^. Left: Read depth at each nucleotide normalized to the number of reads aligning to the 3’-end of 18S rRNA. Nucleotide positions in 18S rRNA are indicated. Above each graph is a schematic of the 18S rRNA and the ITS1 (internal transcribed spacer 1) sequence above their corresponding nucleotide position and read depth. A black line indicates the 3’-end of 18S rRNA. Read depth over the 21 nucleotides at the 3’-end of 18S rRNA (A) and the 20 nucleotides at the 5’ end of ITS1 (B). Right: The fraction of reads miscleaved after each of the final 5 nucleotides of 18S rRNA (A) or after each of the first 5 nucleotides in ITS1 (B) surrounding the canonical 3’-end of 18S rRNA. Data are the average of 8 biological replicates, and error bars indicate standard error of the mean (SEM) (error bars are too small to be seen for many data points). (C) Doubling times of NOY504 cells depleted of endogenous rRNA via growth at 37°C and expressing plasmid-encoded wild type (WT) 18S rRNA (GPD promoter), miscleaved 18S rRNAs, or an empty vector (e.v.). Data are the averages of 14-21 biological replicates, and error bars indicate SEM. N.S., not statistically significant, * p_adj_<0.05, *** p_adj_<0.001, **** p_adj_<0.0001, by one-way ANOVA (Dunnett’s multiple comparisons test). (D) Left: Northern blot of total RNA from cells in panel C. Plasmid-encoded 18S rRNA and 25S rRNA each contain a neutral, unique sequence that is not present in endogenous rRNAs and was specifically detected with a Northern probe (18S Tag and 25S Tag, respectively)^32^. Additional probes were used to detect all 18S or 25S rRNAs using sequences common to the plasmid and endogenous rRNAs (18S and 25S, respectively). All samples were run on the same Northern blot and the order was edited for clarity. Right: Levels of plasmid-encoded 18S rRNAs were normalized to U2 snRNA. Data are the averages of 2-3 biological replicates, and error bars indicate SEM. As previously observed, cells encoding mutant 18S rRNAs are under selective pressure and can undergo homologous recombination with the endogenous 18S rDNA, resulting in a loss of the 18S rRNA tag and the miscleavage phenotype^39^. We have therefore excluded such replicates from quantification. (E) Northern blots of 10-50% sucrose gradients from lysates of cells in panel C. Northern blots were probed for plasmid-encoded 18S and 25S rRNAs (Tag), as well as all 18S and 25S rRNAs. Fraction numbers are listed above the Northern blots. Absorbance profiles shown in Figure S1C. (F) Quantification of the plasmid-encoded 18S rRNA (18S Tag) levels normalized to the plasmid-encoded 25S rRNA (25S Tag) in each polysome fraction (fractions 8-13) from panel E.

### Cell growth is perturbed upon expression of miscleaved rRNAs

Above, we observed that miscleaved 18S rRNAs are rare *in vivo*. However, previous *in vitro* data indicated that Nob1 frequently miscleaves RNA^5,20,33,34^. This discrepancy led us to hypothesize that additional mechanisms might exist *in vivo* to eliminate miscleaved 18S rRNAs. This would be important if miscleaved 18S rRNA have a detrimental effect on cellular fitness. To determine whether ribosomes containing miscleaved 18S rRNA have a negative impact on cell growth, we took advantage of a temperature sensitive *S. cerevisiae* strain (NOY504) containing a deletion of a nonessential subunit of RNA polymerase I, RPA12 (RRN4)^45^. At nonpermissive temperatures, this mutation reduces RNA polymerase I (PolI) complex stability, thus limiting PolI transcription to less than 5% of that at permissive temperature^46^. NOY504 cells were transformed with plasmids encoding an RNA polymerase II promoter-driven 35S rDNA (which encodes the 18S rRNA, 5.8S rRNA, and 25S rRNA)^47^. This plasmid-encoded rRNA can be distinguished from endogenous rRNA by a sequence tag in 18S rRNA (Gal7 promoter constructs) or by sequence tags in both 18S and 25S rRNA (GPD promoter constructs), which are functionally neutral^32,48–50^. Therefore, at the permissive temperature (30°C), cells express both endogenous and plasmid-encoded 18S rRNA, 5.8S rRNA, and 25S rRNA, with endogenous rRNA in vast excess^39^. However, at 37°C, the endogenous rRNA is not transcribed and the cells rely solely on the plasmid-encoded rRNA transcribed by RNA polymerase II (PolII).

To delineate the effects from rRNA miscleavage on cell viability and translation, we encoded truncated 18S rRNA on the plasmid, thereby rendering all plasmid-encoded rRNA “miscleaved” (Figure S1C). We were unable to test the effects of elongated 18S rRNA, miscleaved within ITS1, on cell viability because this required the expression of the rRNAs on two separate plasmids (one containing the miscleaved, elongated 18S rDNA template and a second plasmid containing the 5.8S and 25S rDNAs), and cells expressing rRNA from two plasmids grew too slowly to be measured reliably in our system. However, as indicated above, previous data demonstrate that rRNAs retaining three extra nucleotides do not support cell growth^32^. We first compared the growth of NOY504 cells expressing wild type (WT) or miscleaved, truncated rRNA variants by measuring their doubling times at nonpermissive temperature in a continuous growth assay. While shortening the rRNA by 1 nucleotide does not result in any growth defects, cells expressing rRNAs mimicking miscleavage 2-4 nucleotides upstream of the canonical cleavage site produced 1.5-2 fold slower growth compared to cells expressing WT 18S rRNA, nearly as slow, or as slow as cells lacking plasmid-encoded rRNA entirely (Figure 1C). Importantly, both WT and miscleaved 18S rRNAs produce similar amounts of plasmid-derived tagged 18S rRNA (Figure 1D), demonstrating that the growth defects cannot be explained by reduced amounts of rRNA. Moreover, these miscleaved RNAs are actively translating, as gradient sedimentation shows that the abundance of the 18S-tag relative to the 25S-tag is the same throughout the polysome fractions, if not more abundant within the heavy polysomes (Figure 1E-F and Figure S1D). Thus, miscleaved 18S rRNAs can translate mRNAs but are somehow defective, leading to substantial growth defects.

In addition, miscleaved rRNAs do not accumulate rRNA precursors (Figure 1D), strongly suggesting that Nob1 may only recognize the 3’-A, which is common to all miscleaved substrates and thus lacks strong sequence specificity. This observation, together with the frequent miscleavage observed *in vitro*^5,20,33,34^, raises the question of how the uniformity of the 18S rRNA 3’-end that we observe in WT cells is achieved.

In the above experiments, we utilized a system in which most 18S rRNA is miscleaved. However, as also shown above, in an authentic miscleavage context, cells produce only a small proportion of miscleaved 18S rRNA alongside mostly correctly cleaved 18S rRNAs. To mimic this, we measured the growth of a WT *S. cerevisiae* strain with fully functional RNA polymerase I (BY4741) expressing the same plasmid-encoded WT or truncated, miscleaved 18S rRNAs. Surprisingly, compared to expression of plasmid-encoded WT 18S rRNA, mutant ribosomes containing 18S rRNAs miscleaved 2 or 3 nucleotides upstream of site D caused small, but significant dominant-negative growth defects, despite being vastly outnumbered by endogenous ribosomes. The 18S-4 mutant also caused a slight, although non-significant growth defect (Figure 2A). Thus, in the presence of correctly matured rRNA, miscleaved 18S rRNAs, even in small amounts, perturb the ability of cells to grow, suggesting that translating ribosomes containing miscleaved 18S rRNAs disrupt translation by subunits containing correctly cleaved 18S rRNA.

**Figure 2:**
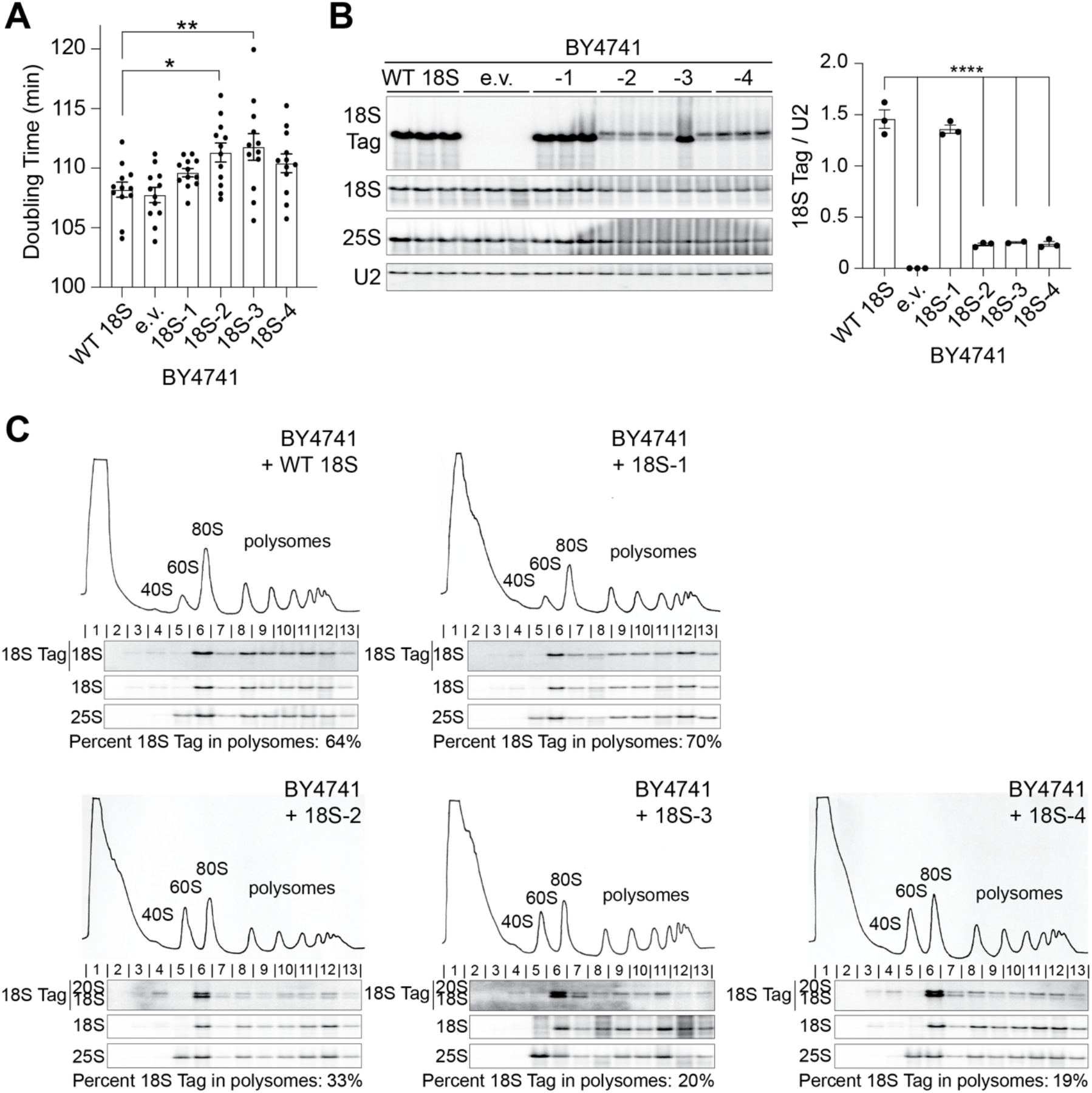
Miscleaved 18S rRNAs perturb the translation of correctly matured 40S. (A) Doubling times of BY4741 cells expressing both endogenous rRNAs and plasmid-encoded 18S rRNAs (Gal7 promoter) or an empty vector (e.v.) grown at 30°C. Data are the averages of 12 biological replicates, and error bars indicate SEM. *p_adj_<0.05, **p_adj_<0.01 by one-way ANOVA (Dunnett’s multiple comparisons test). All other differences in doubling time were not statistically significant. (B) Left: Northern blot of total RNA from cells in panel A. Right: Plasmid-encoded 18S rRNA accumulation was normalized to U2 snRNA. Data are the averages of 2-3 biological replicates, and error bars indicate SEM. The second replicate of 18S -3 was excluded from analysis as explained in the legend of Figure 1. **** p_adj_<0.0001, by one-way ANOVA (Dunnett’s multiple comparisons test). The difference in 18S Tag/U2 accumulation between WT 18S and 18S-1 was not statistically significant. (C) 10-50% sucrose gradients of lysates from cells in panel A. Below the absorbance profile at 254 nm are Northern blots of the plasmid-encoded 18S rRNA (Tag), as well as total 18S and 25S rRNAs. The percent of plasmid-encoded 18S rRNA or 20S pre-rRNA in polysomes (fractions 8-13) compared with total plasmid-encoded 18S rRNA or 20S pre-rRNA was calculated and is listed below the Northern blot.

### Miscleaved rRNAs are targeted for collision-mediated decay

Given the dominant-negative growth defect of miscleaved 18S rRNAs, despite their low abundance in BY4741 cells, the simplest model is that the miscleaved ribosomes perturb translation of all ribosomes by slowing or stalling on mRNA. Slower or stalled ribosomes lead to ribosome collisions, which result in the decay of the translated mRNA via no-go decay (NGD), the degradation of the nascent peptide chain through ribosome-associated quality control (RQC), and to 18S non-functional rRNA decay (NRD), which targets non-functional 18S rRNAs for degradation^35,36^. Thus, if miscleaved rRNAs lead to ribosome collisions with correctly matured 40S leading to degradation of the miscleaved compromised 18S rRNA, a prediction would be that in WT yeast (which contain mostly correctly matured 18S rRNA) the miscleaved rRNAs are reduced in their abundance relative to functional WT rRNAs. To test this prediction, we carried out Northern blotting of total RNA from WT BY4741 cells supplemented with plasmids encoding either WT or miscleaved rRNAs. In these cells, we observe a ∼5 fold reduction in 18S rRNAs miscleaved by 2-4 nucleotides compared to WT 18S rRNA (Figure 2B), even though no reduction in 18S rRNA from the same constructs was observed when the miscleaved rRNAs were the only expressed RNAs (Figure 1D). The reduction of miscleaved 18S rRNA only in cells with correctly assembled endogenous ribosomes is consistent with ribosome collisions leading to degradation of the miscleaved 18S rRNA. When all ribosomes are equally defective, and therefore move at the same (slow) speed, as in the NOY504 cells, collisions are less likely, as previously shown when some ribosomes are slowed by binding of translation inhibitors like cycloheximide ^51^.

To further confirm that miscleaved 18S rRNA are cleared via ribosome collisions when wt ribosomes are present, we used gradient sedimentation to monitor where plasmid-encoded 18S rRNAs sediment. If the ribosomes containing miscleaved rRNAs collide with endogenous, correctly matured 40S leading to decay of the defective ribosomes, one would predict that they are selectively depleted from heavy polysomes, where the density of ribosomes is higher and the chances for collisions are therefore larger^52^. Indeed, while WT 18S and 18S-1 rRNAs are equally abundant throughout the polysome fractions, miscleaved 18S-2, 18S-3, and 18S-4 rRNAs are more abundant in the monosome and light polysome fractions than in the heavy polysome fractions, consistent with miscleaved 18S rRNA causing ribosome collisions and being degraded (Figure 2C). Importantly, as shown before, absence of the miscleaved rRNAs in the polysomes is not a reflection of their inability to translate, as they are recruited to the polysomes and stable, when they are the only 18S rRNAs in the cell (Figure 1E-F), rather, it is indicative of the miscleaved 18S rRNA-containing ribosomes having a translation elongation defect, leading to collisions by WT ribosomes.

To further confirm that miscleaved 18S rRNAs are being turned over in response to interactions with fully functional endogenous ribosomes, we tracked the abundance of miscleaved 18S rRNAs over time as miscleaved 18S rRNA transcription was activated or as endogenous ribosomes were depleted (Figure 3A-C). If miscleaved ribosomes translate more slowly, leading to collisions by endogenous ribosomes that ultimately lead to their decay, then under conditions when endogenous ribosomes are being depleted, the frequency of collision events and subsequent rRNA decay should decrease over time, thus stabilizing miscleaved 18S rRNA. Indeed, while initially miscleaved 18S rRNA (closed symbols) is less abundant than control WT rRNA (open symbols), 3-4 cell doublings after rRNA transcription shutoff (by shifting the NOY504 strain from 30 °C to 37 °C), which reduces the amount of endogenous rRNA by 8-16 fold, the abundance of constitutively expressed miscleaved 18S rRNA, but not that of WT control 18S rRNA, is increased. This ultimately leads to equal abundance of WT and miscleaved 18S rRNA (left and middle panel in Figure 3A-B and black and green symbols in Figure 3C), as observed in steady-state experiments (Figure 1D). Moreover, the delay in accumulation of miscleaved 18S rRNA is about one doubling shorter for cells with plasmids driven by the very strong Gal7 promoter, than the weaker GPD promoter (Figure 3C, blue and black closed symbols), as expected because more plasmid-derived rRNA is produced from the Gal7 promoter. Finally, we note that induction of the plasmid-encoded miscleaved 18S rRNA (by a change in the sugar source from raffinose to galactose) accumulates less miscleaved 18S rRNA in cells where endogenous ribosomes are not depleted relative to cells where endogenous ribosomes are depleted (Figure 3B, open symbols in Figure 3C). In contrast, WT 18S rRNA accumulates to the same extent in all cells, demonstrating that this is not a transcriptional effect (Figure 3A, closed symbols in Figure 3C). Furthermore, the kinetics of accumulation are slower with the miscleaved rRNA relative to WT rRNA (blue symbols in Figure 3C). Together, these data strongly support the model that miscleaved 18S rRNAs are degraded by a mechanism that is sensitive to the presence and concentration of 40S subunits containing endogenous, correctly cleaved 18S rRNA.

**Figure 3:**
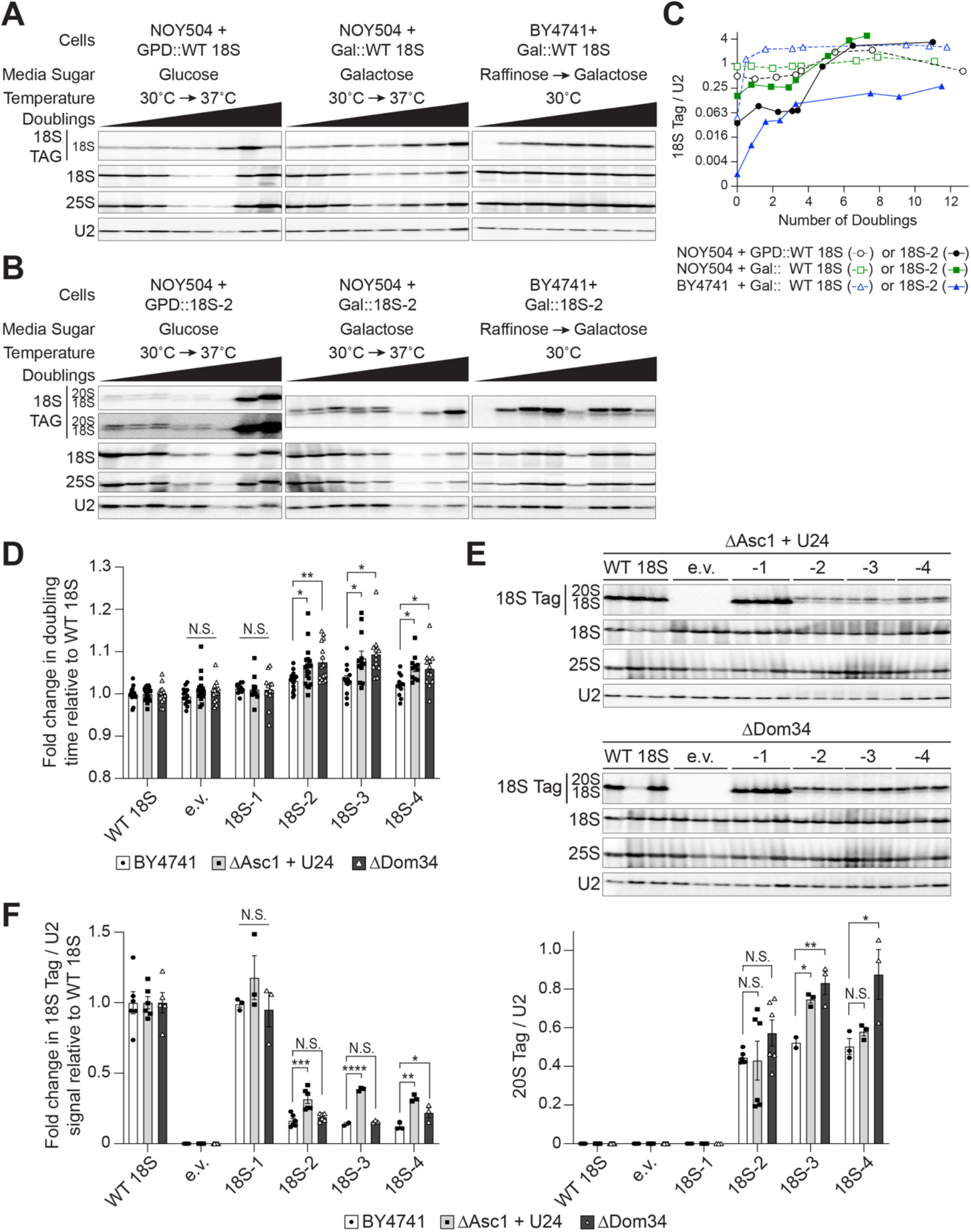
Miscleaved 18S rRNAs lead to ribosome collisions and are degraded. (A-C) Total RNA Northern blots of plasmid-encoded WT 18S (A) or miscleaved 18S-2 rRNA (B) over time. In each panel, cells were first grown to mid-log phase in one condition (first time point), and then switched to a second growth condition (subsequent time points taken over time). In the first two panels, NOY504 cells expressed 18S rRNA from either a GPD promoter grown in glucose dropout media (left panel) or from a Gal7 promoter grown in galactose dropout media (middle panel). The cells were first grown at 30°C to express both endogenous and plasmid rRNAs, and then switched to 37°C where only plasmid rRNAs were transcribed. Finally, in the right panels, BY4741 cells expressing 18S rRNA from a Gal7 promoter were grown at 30°C first in raffinose dropout media (endogenous rRNA was transcribed with minimal plasmid rRNA transcription) and switched to galactose dropout media to induce plasmid rRNA transcription in the presence of endogenous rRNA expression. (C) Plasmid-encoded WT 18S (open circles) or miscleaved 18S-2 rRNA (closed circles) levels were normalized to U2 snRNA at each time point and plotted against the number of cell doublings following the switch in growth conditions (either temperature or sugar). The first time point, before the switch in growth conditions, is indicated as 0 doublings. (D) Changes in doubling time of wild type cells (BY4741), cells lacking Asc1 (ΔAsc1), or cells lacking Dom34 (ΔDom34), each supplemented with plasmids encoding WT 18S, an empty vector, or miscleaved 18S rRNAs under a Gal7 promoter grown at 30°C. ΔAsc1 cells were supplemented with a plasmid encoding U24 snRNA, normally encoded in the *ASC1* intron. Doubling times for BY4741 cells include those from Figure 2A and additional replicates. Doubling times were normalized to WT 18S for each cell background (fold change = 1). Data are the averages of 12-19 biological replicates, and error bars indicate SEM. N.S. not statistically significant, *p_adj_<0.05, **p_adj_<0.01, by one-way ANOVA (Dunnett’s multiple comparisons test) compared to wild type cells for each 18S rRNA variant. (E) Northern blots of total RNA from cells in panel D. (F) Quantification of Northern blots in panel E, Figure 2B, and additional replicates not shown. Plasmid-encoded 18S rRNA or 20S pre-rRNA were normalized to U2 snRNA. 18S/U2 ratios from cells expressing 18S-2 were normalized to the 18S/U2 ratios from cells expressing WT 18S for each cell background (fold change = 1). Data are the averages of 2-6 biological replicates, and error bars indicate SEM. N.S. not statistically significant, *p_adj_<0.05, **p_adj_<0.01, ***p_adj_<0.001, ****p_adj_<0.0001, by one-way ANOVA (Dunnett’s multiple comparison’s test) compared to BY4741 for each 18S rRNA variant.

While the data above are consistent with degradation of miscleaved 18S rRNA induced by ribosome collisions, they could also be consistent with other yet-unknown pathways. Thus, we sought to provide additional evidence linking the decay of these miscleaved ribosomes to known factors that recognize the collided ribosomes and target them for disassembly and decay. Collided ribosomes form a unique interface involving the 40S RP Asc1, which is recognized by the E3 ubiquitin ligases Mag2 and Hel2 to ubiquitinate Rps3 (uS3). This leads to subunit dissociation either by the RNA helicase Slh1 or by Dom34:Hbs1. Degradation of the non-functional 18S rRNA relies on the cytoplasmic exosome recruitment factor Ski7 and the exonuclease Xrn1^40–42,53–57^. If correctly cleaved endogenous ribosomes collide with miscleaved 18S rRNAs leading to their decay, then deleting these factors should render cells defective in clearing miscleaved 18S rRNA-containing ribosomes, thereby exacerbating their dominant-negative growth defect. By analyzing the doubling time of yeast in a continuous growth assay, we observed that deleting Dom34, Asc1, or Hel2 sensitizes cells to miscleaved 18S rRNAs, while the deletion of Xrn1 partially rescued the growth defect of miscleaved 18S rRNA (Figure 3D and Figure S2A). As expected if these proteins play a role in degradation of miscleaved 18S rRNA, deleting Asc1, Hel2, or Xrn1 stabilized plasmid-derived miscleaved 18S rRNA, but not the 20S pre-rRNA (Figure 3E-F and Figure S2B), as expected because most 20S pre-rRNAs are not translating^3,58^. Deletion of Dom34 also stabilized the 4-nucleotide truncated, miscleaved 18S rRNA, but not the other shortened rRNAs. Moreover, Dom34 deletion significantly stabilized the plasmid-encoded immature 20S pre-rRNA (Figure 3E-F). Both observations likely explained by the role Dom34 plays during pre-40S assembly in separating 80S-like ribosomes, resulting in 20S pre-rRNA accumulation and 18S rRNA depletion upon Dom34 deletion^3^, which would partially mask the effect of collisions. In summary, our data strongly support a role for ribosome collisions in the decay of 3’-end miscleaved 18S rRNAs, specifically involving Asc1, Hel2, Dom34 and Xrn1 and the presence of functional mature ribosomes.

### Bypassing Rio1 stabilizes miscleaved 18S rRNAs

Above, we have shown that Nob1 has limited sequence specificity as it cleaves the truncated 18S rRNA mutant constructs, which present an incorrect cleavage site (Figure 2B and S1B), consistent with observations that demonstrate miscleavage *in vitro*^5,20,33,34^. Yet, we have also shown that the majority of 18S rRNA in WT cells is cleaved correctly and that maintaining accurate cleavage is important because miscleaved rRNAs, even in small amounts, induce collisions that perturb translation globally (Figure 2-3 and Figure S2). Together, these observations raise the question: how do cells maintain fidelity given that it is not entrusted to Nob1? Clearly, decay of miscleaved rRNAs after maturation is part of the answer as demonstrated above. This pathway, if overloaded, induces a cellular stress response^59^. Moreover, miscleaved 18S rRNAs are dominant negative (Figure 2A) even if they make up only ∼1% of total cellular 18S rRNA^39^. Thus, reducing the amount of miscleaved 18S rRNA that enters the translating pool is critical.

We therefore hypothesized that correct 18S rRNA cleavage was monitored by the Rio1-mediated checkpoint that prevents release of immature rRNA into the translating pool^7^. This checkpoint is established as Nob1 and Pno1 cooperate to prevent pre-40S ribosomes from initiating translation prematurely. Nob1 and Pno1 are released by the kinase Rio1, but only after Nob1 has cleaved 18S rRNA^7,26^. Thus, this QC checkpoint ensures that only mature, 18S rRNA-containing ribosomes engage in translation^7^. Importantly, if Rio1 required cleavage accuracy, this checkpoint could also allow for monitoring of correct cleavage of the 3’-ends of 18S rRNA. If this hypothesis is correct, we would expect miscleaved 18S rRNAs to be more abundant in cells that bypass this QC step.

To test this prediction, we first introduced a mutation in Pno1 that bypasses this QC step. Pno1-KKKF (K208E/K211E/K213E/F214A) disrupts Pno1’s contact at the 3’-end of 18S rRNA^26,29^ (Figure S3A) thereby weakening the binding of Pno1 to the pre-40S ribosome^60^, resulting in Rio1-independent release of Pno1 (and Nob1 which is weakly bound in the absence of Pno1)^7^. We performed 3’-RACE-sequencing of 18S rRNA from 40S ribosomes purified from cells depleted of endogenous Pno1 and Dim1 and expressing WT Pno1 or Pno1-KKKF and the dimethylation-deficient Dim1-E85A^44^. 96-99% of the 2.8-4.5 million reads per sample we obtained map to the 3’-end of 18S rRNA. 2.5% of 18S rRNAs are miscleaved in cells expressing Pno1-KKKF, compared to only 1.9% miscleavage in wild type cells (Figure 4A and S3B). Specifically, products miscleaved upstream of the correct 18S rRNA 3’-end are more abundant in cells expressing Pno1-KKKF compared to cells expressing WT Pno1 (Figure 4A and S3C). In agreement with no growth defect caused by 18S-1 (Figures 1C and 2A), miscleavage at 18S-1 was not significantly different in cells expressing WT Pno1 or Pno1-KKKF. Miscleavage in ITS1 downstream of the 18S rRNA 3’-end is also not significantly altered upon Pno1-KKKF expression (Figure S3B-C). Importantly, the distribution of miscleavage events is the same in WT Pno1 and Pno1-KKKF cells (Figure 4A), demonstrating that changes in the overall rate of miscleavage were not due to changes in cleavage site recognition by Nob1 and indicating that Nob1 activity remained unperturbed. Thus, bypassing Rio1 increases the abundance of miscleaved 18S rRNAs. Importantly, this is specific to 18S rRNA 3’-end cleavage, as bypassing Rio1 does not affect miscleavage at the 3’-end of 25S rRNA (Figure S3D).

**Figure 4:**
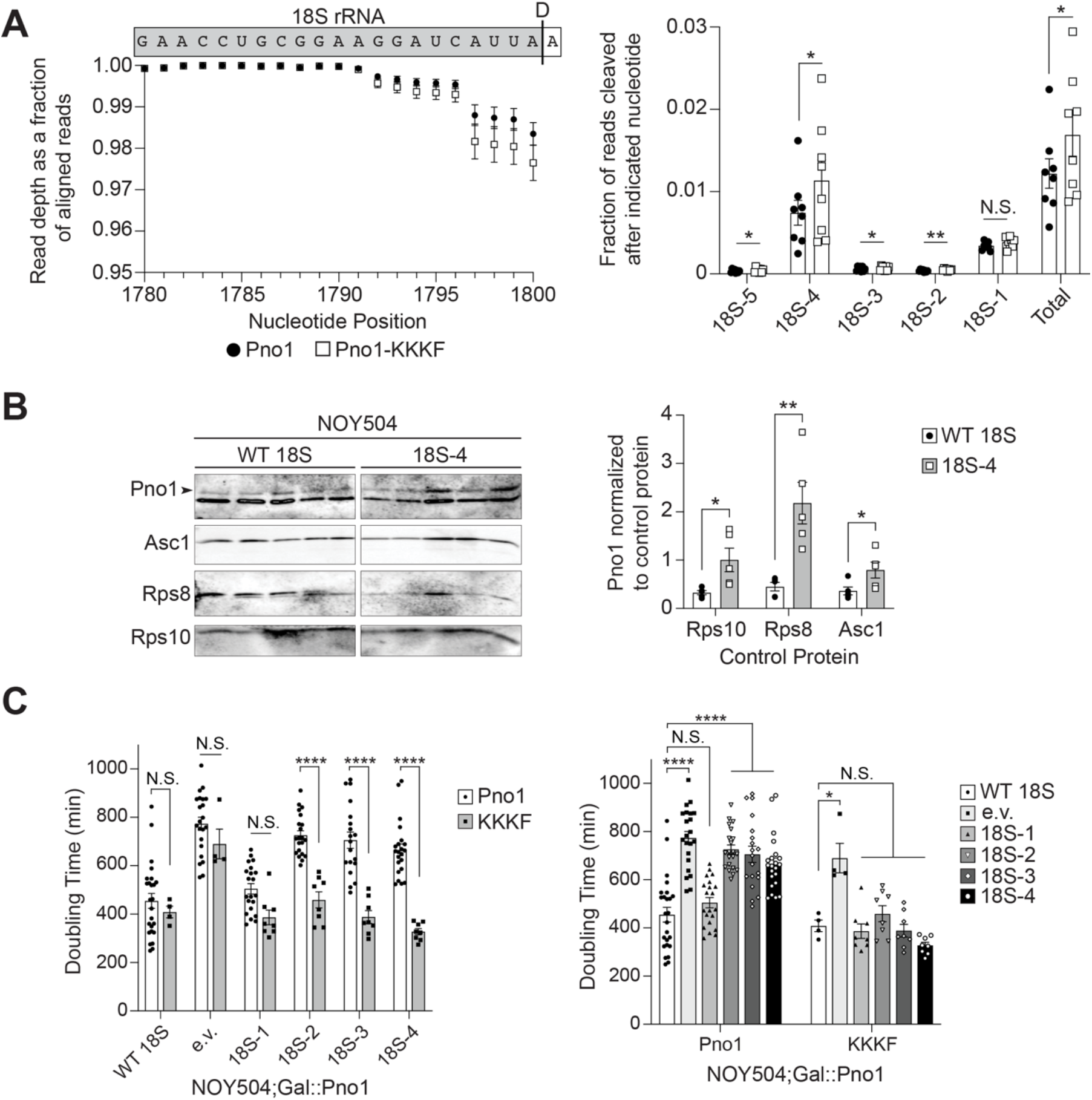
Bypassing Rio1 stabilizes miscleaved 18S rRNAs. (A) 3’-RACE-sequencing of 18S rRNAs extracted from 40S ribosomal subunits purified from Gal::Pno1; Gal::Dim1 cells depleted of endogenous Pno1 and Dim1 by growth in glucose and supplemented with plasmids expressing Dim1-E85A and either Pno1 or Pno1-KKKF (K208E/K211E/K213E/F214A), which bypasses the Rio1-mediated QC step during pre-40S ribosome assembly^7^. The wt Pno1 data is the same as in Figure 1A-B. Left: Read depth at each nucleotide normalized to the number of reads aligning to the 3’-end of 18S rRNA, upstream of the cleavage site. Above the graph is a schematic of the 18S rRNA and the ITS1 sequence above their corresponding nucleotide position and read depth. A black line indicates the D cleavage site that forms the 3’-end of 18S rRNA. Right: The fraction of reads miscleaved after each of the final 5 nucleotides of 18S rRNA. “Total” represents the cumulative miscleavage from 18S-5 to 18S-1. Data are the average of 8 biological replicates, and error bars indicate SEM (error bars are too small to be seen for many data points). N.S. not statistically significant, *p<0.05, **p<0.01, by ratio paired t-test comparing miscleavage in Pno1 and Pno1-KKKF for each nucleotide. Pno1 and Pno1-KKKF samples grown and analyzed on the same day were considered paired replicates. (B) Left: Western blot of MS2-tagged RNA affinity-purified ribosomes containing WT or miscleaved 18S-4 rRNA. Both contain an internal MS2 RNA hairpin and are transcribed from a plasmid (GPD promoter). NOY504 cells were grown at 37°C. The arrowhead notes the upper band corresponding to Pno1. Right: Quantification of Pno1-bound ribosomes normalized to either Rps10, Rps8, or Asc1 control proteins as indicated. Data are the average of 5 biological replicates, and error bars indicate SEM. *p<0.05, **p<0.01, by unpaired t-test comparing Pno1 abundance on WT 18S to 18S-4 ribosomes for each control protein. (C) Doubling times of NOY504;Gal::Pno1 cells grown at 37°C, depleted of endogenous Pno1 by growth in glucose, and expressing either Pno1 or Pno1-KKKF and either WT 18S, an empty vector (e.v.), or miscleaved 18S rRNAs from a GPD promoter. Data are the averages of 4-25 biological replicates, and error bars indicate SEM. N.S. not statistically significant, *p_adj_<0.05, ****p_adj_<0.0001, by two-way ANOVA (Tukey’s multiple comparisons test). The graph on the right is the same data as the graph on the left, presented in a different order for clarity in representing statistical comparisons.

If the Rio1-mediated checkpoint is involved in surveillance of 18S rRNA miscleavage, as suggested by the data above, then we would expect that miscleaved 18S rRNA that escape into the translating pool retain Pno1. To test this prediction and further confirm a role for Rio1 in monitoring cleavage accuracy, we grew NOY504 cells at 37°C expressing either WT 18S rRNA or miscleaved 18S-4 rRNA containing MS2 hairpins loops and purified plasmid-derived ribosomes via MS2-tagged RNA affinity purification. Indeed, western blots indicated that relative to three RPs, Pno1 is significantly more abundant on the ribosomes containing -4 miscleaved 18S rRNA than their WT counterparts (Figure 4B). Unfortunately, we were unable to measure whether Nob1 is also retained on miscleaved ribosomes because the bait protein for purification, MS2-MBP, co-migrates with Nob1 on a Western blot, and cross-reacts with the Nob1 antibody.

To further validate that miscleaved 18S rRNA-containing ribosomes retain Pno1, we tested whether weakly binding Pno1 mutants could partially rescue the slow growth phenotype of miscleaved 18S rRNAs. Pno1-KKKF rescues the growth defect of cells expressing miscleaved 18S rRNAs truncated by 2-4 nucleotides, with no significant effect on the growth of cells expressing miscleaved 18S rRNAs truncated by a single nucleotide (Figure 4C). Therefore, Pno1 cannot be removed from these ribosomes, showing that miscleaved ribosomes fail the QC checkpoint mediated by Rio1.

### Rio1 binds miscleaved RNAs more weakly

We have previously observed that overexpression of Rio1 in the presence of an inactive Nob1 mutant releases 20S pre-rRNA-containing ribosomes into the translating pool^7^, indicating that Rio1’s selectivity arises from differences in its affinity for 20S or 18S-containing ribosomes. We therefore wanted to test directly whether Rio1 was able to distinguish correctly from incorrectly cleaved rRNAs based on differential binding ability. To test this hypothesis, we used a previously described quantitative *in vitro* RNA binding assay^19^ to measure the binding of Rio1 to *in vitro* transcribed RNA mimics of 18S rRNAs with variable 3’-ends. These mimics contained the 3’-end of 18S rRNA, starting at h44 and ending at either the correct 3’-end, 3 nucleotides further, or 4 nucleotides short. The RNAs were transcribed from PCR products generated with primers containing two 2’-O-methylated RNA nucleotides to ensure the precision of the 3’-end^61^, folded, and then incubated with Rio1. Rio1-bound and free RNAs were separated by native PAGE gels. Comparing the binding of Rio1 to these 18S rRNA mimics, our data show 3-fold stronger Rio1 binding to the RNA mimic of the correctly cleaved 18S rRNA (H44-D) compared to the RNA mimic of the truncated, miscleaved 18S rRNA (H44-D-4). Rio1 also binds the correctly cleaved 18S rRNA slightly (1.4-fold) stronger than its binding to the elongated, miscleaved 18S rRNA (H44-D+3, Figure 5A-B). The preferential binding to accurately processed 18S rRNA suggests that Rio1 directly senses the sequence and/or length at the 3’-end of 18S rRNA.

**Figure 5:**
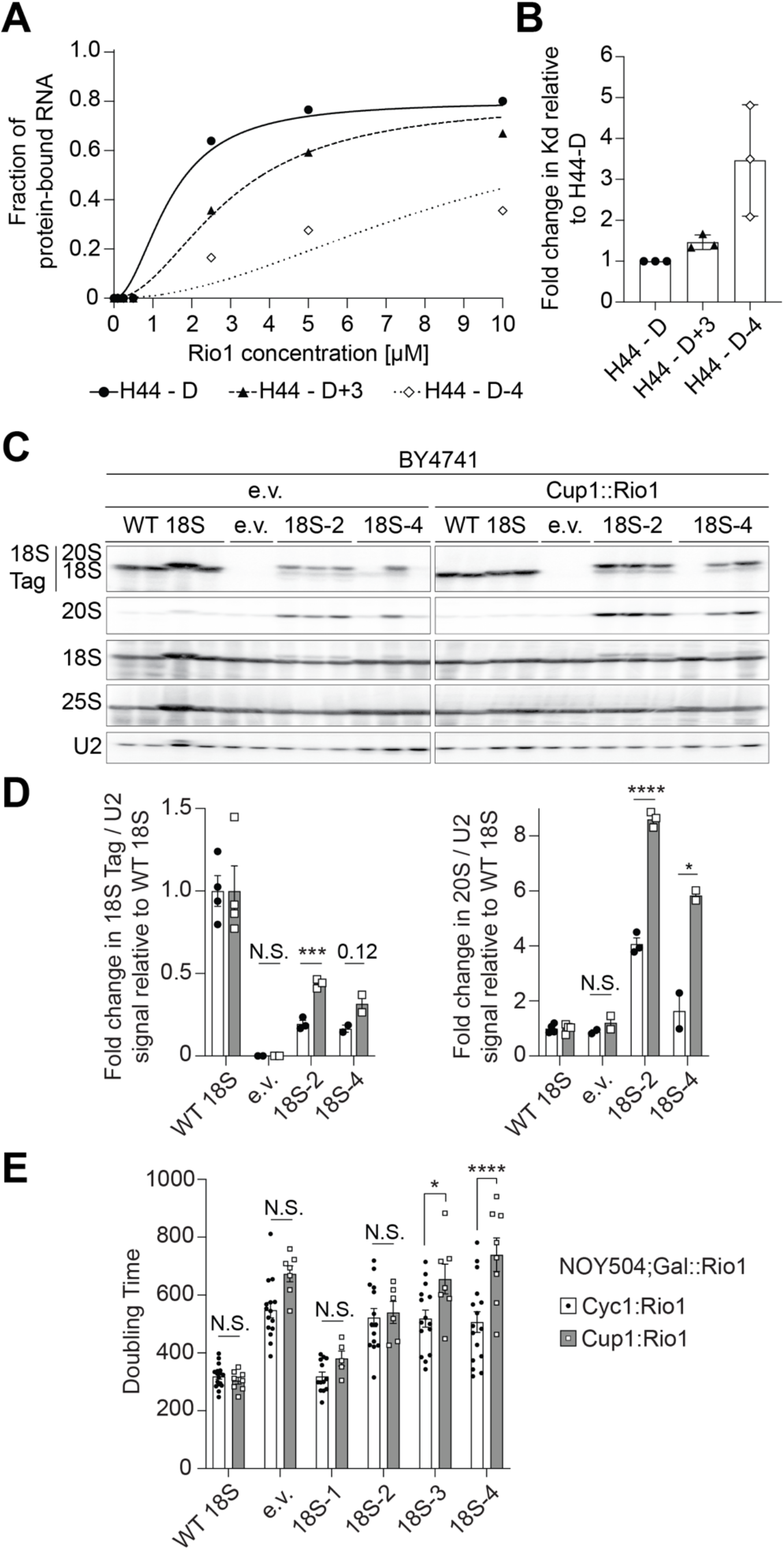
Rio1 monitors 18S rRNA cleavage during pre-40S assembly QC. (A) Representative RNA binding assay with *in vitro* transcribed H44-D (18S rRNA mimic, black circles), H44-D+3 (+3 nt, long miscleaved 18S rRNA mimic, black triangles), and H44-D-4 (−4 nt, short miscleaved 18S rRNA mimic, white diamonds) RNAs and recombinant Rio1. Three independent experiments resulted in K*_d_* = 2.3 +/- 1.0 µM^2^ for Rio1 binding H44-D, K*_d_* = 3.3 +/- 1.2 µM^2^ for Rio1 binding H44-D+3, K*_d_* = 7.5 +/- 1.9 µM^2^ for Rio1 binding H44-D-4. (B) To account for day-to-day variations, K*_d_* values of Rio1 binding each RNA mimic from panel B were normalized to the K*_d_* for Rio1 binding H44-D on each gel (fold change = 1). (C) Total RNA Northern blot from BY4741 cells expressing excess Rio1 under a Cup1 promoter or only endogenous Rio1 (e.v., empty vector) and either WT 18S, an empty vector, or miscleaved 18S rRNAs from a Gal7 promoter. (D) Quantification of Northern blots in panel C. Plasmid-encoded 18S rRNA or total 20S pre-rRNA was normalized to U2 snRNA. 18S Tag/U2 ratios from cells expressing miscleaved 18S rRNAs were normalized to the 18S Tag/U2 ratios from cells expressing WT 18S for each cell background (fold change = 1). Data are the averages of 2-4 biological replicates, and error bars indicate SEM. N.S. not statistically significant, *p<0.05, ***p<0.001, ****p<0.0001, by unpaired t-test for each 18S rRNA variant. (E) Changes in doubling time of NOY504;Gal::Rio1 cells grown at 37°C, depleted of endogenous Rio1 by growth in glucose, expressing Rio1 either under a Cyc1 promoter (near endogenous expression level) or a Cup1 promoter (high expression level) and either WT 18S, an empty vector, or miscleaved 18S rRNAs from a GPD promoter. Cells expressing Cup1::Rio1 were grown in media supplemented with 10 µM CuSO_4_ to activate the Cup1 promoter. Data are the averages of 5-17 biological replicates, and error bars indicate SEM. N.S. not statistically significant, *p<0.05, ****p<0.0001, by unpaired t-test for each 18S rRNA variant.

To further test if weakened binding to miscleaved RNAs *in vivo* could account for Rio1’s ability to QC cleavage accuracy, we tested whether we could rescue Rio1’s weakened binding to miscleaved 18S rRNA-containing ribosomes by overexpressing Rio1. If so, then we expect that miscleaved 18S rRNAs should become more abundant. Indeed, in cells expressing endogenous rRNA (BY4741) and excess Rio1 (Cup1 promoter, Figure S4), we observed increased accumulation of miscleaved 18S rRNA (Figure 5C-D). We also saw an increase in 20S pre-rRNA abundance, as the extra Rio1 also releases Nob1 and Pno1 prematurely from un-cleaved ribosomes^7^. Next, we grew NOY504 cells at 37°C to express only plasmid-encoded 18S rRNAs and depleted endogenous Rio1 under a galactose-inducible/glucose-repressible promoter by growth in glucose. Rio1 was then expressed from a plasmid, either near endogenous levels under the Cyc1 promoter or overexpressed under the copper inducible Cup1 promoter (Figure S4). As expected from an increase in miscleaved 18S rRNA-containing ribosomes, excess Rio1 caused a significant additional growth defect in cells relying solely on ribosomes containing miscleaved 18S rRNAs (Figure 5E). Altogether, these data support a model in which Rio1 monitors the precise cleavage of the 18S rRNA 3’-end during ribosome maturation, marking correctly processed ribosomes by the removal of Nob1 and Pno1 from the ribosome.

## Discussion

### Rio1 monitors 18S rRNA cleavage accuracy during ribosome assembly

In this work, we expand our understanding of the role Rio1 plays in ribosome assembly and show that Rio1 helps ensure that 18S rRNA is cleaved at the correct site. Previously, we demonstrated that Nob1 and Pno1 establish a quality control checkpoint wherein immature ribosomes containing 20S pre-rRNA are prevented from recruiting mRNA by Nob1. Rio1 removes Nob1 and Pno1 in an ATP dependent manner from nascent ribosomes after Nob1 cleaves 18S rRNA, thus licensing only mature ribosomes to recruit and translate mRNA^7^ (Figure 6A).

**Figure 6:**
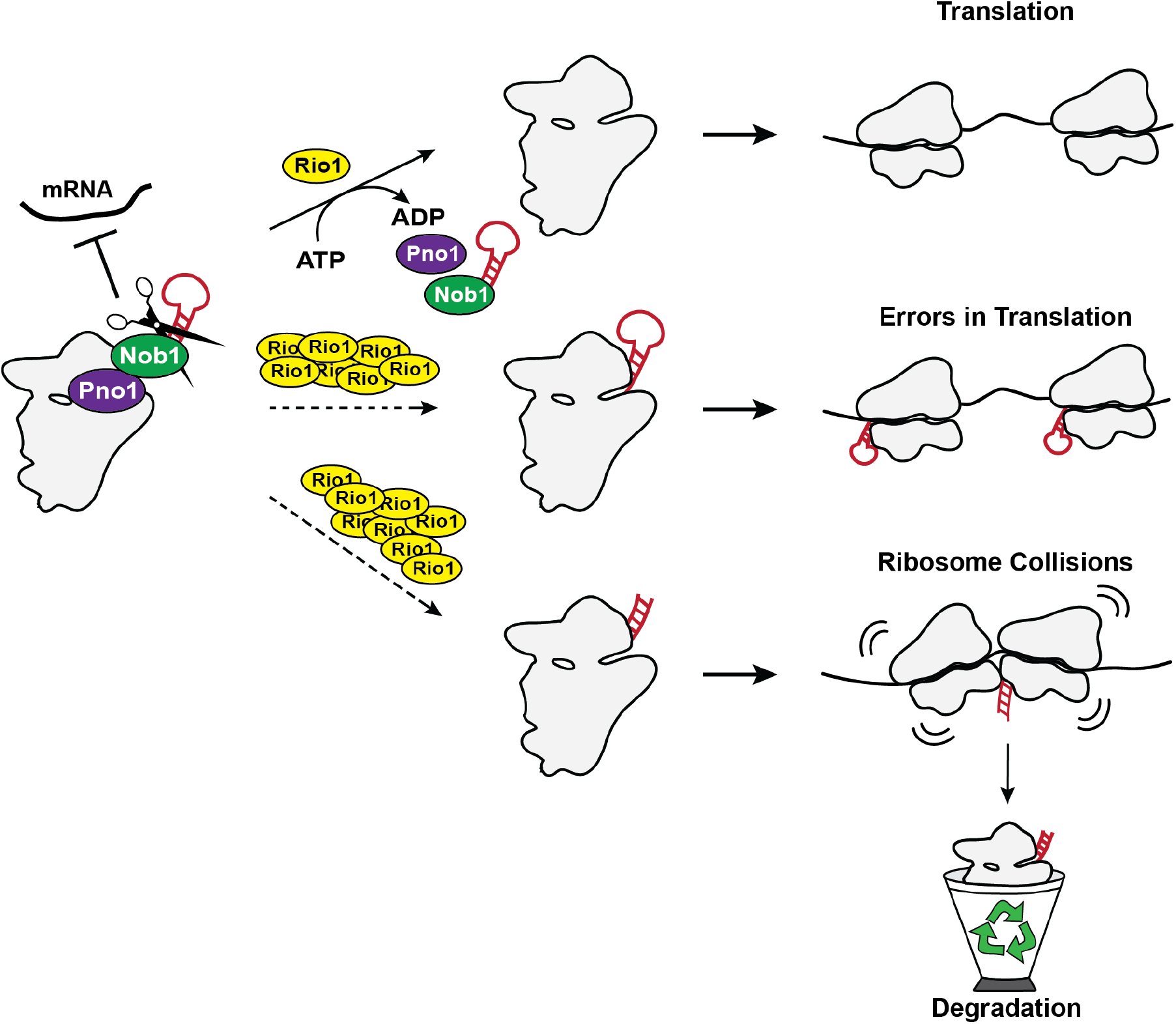
Schematic of the role Rio1 plays in monitoring precise cleavage of the 3’-end of 18S rRNA. Nob1 and Pno1 establish a QC checkpoint in which Nob1 prevents mRNA recruitment to immature ribosomes containing 20S pre-rRNA. (Top) Once Nob1 cleaves the 18S rRNA to form the correct 3’-end, Rio1 releases Nob1 and Pno1 from the nascent ribosome, which can now recruit mRNA^7^. Disrupting the Rio1-mediated QC checkpoint by overexpression of Rio1 (shown here) or mutations in Pno1 that weaken Pno1 binding leads to release of immature ribosomes, which produce translational errors ^7^ (middle), or ribosomes containing miscleaved 18S rRNA (bottom), which lead to collisions with correctly matured ribosomes.

In the current work, we show that Nob1 can miscleave its substrate *in vivo*, confirming earlier observations that purified recombinant Nob1 miscleaves RNA *in vitro*^5,20,33,34^. Miscleaved rRNAs are scarce *in vivo* but become more abundant upon bypass of the Rio1-mediated QC step (Figures 1A-B and 4A-B). Thus, our data support a model in which Rio1 monitors for correct cleavage, restricting miscleaved ribosomes from entering the translating pool. *In vitro* RNA binding data show that Rio1 binds correctly cleaved RNA more strongly than miscleaved RNA (Figure 5A-B) and overexpression of Rio1 increases the amount of miscleaved RNA *in vivo* (Figure 5C-D). Thus Rio1’s ability to monitor correct rRNA cleavage reflects differences in rRNA binding affinity. The importance of this QC step is underlined by our observation that miscleaved rRNAs, even in small quantities, disturb translation.

### Miscleaved 18S rRNAs are more frequently truncated

Through our sequencing analysis, we discovered that *in vivo*, miscleaved 18S rRNAs were truncated within 18S rRNA 10-fold more frequently than they were elongated and cleaved within ITS1 (Figure 1A-B). This is different than what was observed *in vitro*, where Nob1 miscleaves both 3’ and 5’ of the canonical 18S rRNA cleavage site^5,20,33,34^. To reconcile these observations, we consider the differences of Nob1-mediated rRNA cleavage *in vitro* and *in vivo*. *In vitro*, radiolabeling of RNAs allows detection of the elongated and truncated 18S rRNAs. Meanwhile, *in vivo*, if Nob1 miscleaves 18S rRNA 3’ to its cleavage site, the elongated 18S rRNA retains Nob1, preventing these miscleaved ribosomes from entering the translating pool^7^. Nob1 then has a second chance to cleave at the correct nucleotide, thereby correcting its previous mistake. However, if Nob1 truncates 18S rRNA by miscleaving 5’ to its cleavage site within mature 18S rRNA, the Nob1 binding site on the rRNA will be removed leading to Nob1 dissociation and releasing the truncated, miscleaved rRNA-containing ribosomes into the translating pool, where they can be detected in our sequencing assay. Thus, we postulate that the differential abundance of the elongated and shortened miscleavage products reflect Nob1 being able to cleave elongated 18S rRNAs again, rather than the ability of Nob1 to effectively discriminate against 3’-elongated sequences.

Intriguingly, Nob1 appears to prefer cleaving its RNA substrate 5’ to an adenosine, as the most abundant miscleavage events *in vivo* are all followed by an adenosine (Figure 1A-B). Moreover, miscleavage of rRNA substrates by Nob1 *in vitro* also occurs upstream of an adenosine^5,20,34^. Furthermore, while Nob1 is conserved in most archaea, the sequence at the 3’-end of the small subunit rRNA is not. Even so, in the archaea *Pyrococcus horikoshii*, Nob1 cleaves 5’ to an adenosine to form the canonical 16S rRNA 3’-end^34^.

### Monitoring rRNA cleavage

Our data demonstrate the importance of forming the canonical 3’-end of 18S rRNA. The 3’-end of 18S rRNA is highly conserved from yeast to humans (Figure S1A) and our data show that miscleaved 18S rRNAs provide a 1.4- to 1.8-fold growth defect (Figure 1C). While Nob1 can miscleave its rRNA substrate *in vitro* and *in vivo*, we observe very little miscleaved rRNA in cells with functional QC mechanisms. While we cannot rule out that Pno1 remaining on these pre-40S ribosomes contributes to the growth defect, as it blocks binding of the essential RP Rps26^26,28^, the observation that excess Rio1 removes Nob1 and Pno1 from otherwise stalled substrates^7^, but exacerbates the growth defects (Figure 5E) argues against this. Either way, Pno1 bound to miscleaved ribosomes (Figure 4B) is a consequence of the failed QC step in which Rio1 does not remove Nob1 or Pno1 from miscleaved ribosomes. This supports our conclusion that cleavage *accuracy* at the 3’-end of 18S rRNA is quality controlled, and bypass of this QC step leads to nonfunctional ribosomes in the translating pool, where they disrupt translation by correctly processed ribosomes (Figure 6B).

### Rio1 concentration is critical for 18S rRNA QC

Our current work along with previous observations indicates that the concentration of Rio1 is critical for monitoring correct rRNA cleavage and maintaining this QC step. Rio1 has a lower binding affinity for miscleaved rRNAs than correctly cleaved rRNAs *in vitro* (Figure 5A-B). Thus, increasing the amount of available Rio1 leads to Rio1 binding to non-optimal substrates, releasing immature and miscleaved ribosomes into the translating pool (Figure 5C-D). Therefore, the stringency of this QC step, the gate separating late-stage immature pre-40S and mature 40S ribosomes in the cytoplasm, relies on the concentration of Rio1 relative to that of the nascent ribosome pool. Interestingly, whole-genome sequencing of cancer cells revealed that RIOK1, human Rio1, is frequently amplified in cancer: 7% of ovarian tumors, 3.6% of melanomas, 4.2% of mature B-cell neoplasms, 3.8% of ocular melanomas, 3.4% of bladder urothelial carcinomas, 2.7% of liver hepatocellular carcinomas, 2.8% of cholangiocarcinomas, and 2.3% of mesotheliomas (The Cancer Genome Atlas (TCGA): https://www.cancer.gov/tcga, https://www.cbioportal.org). Moreover, RNAseq data show that relative to the mRNAs encoding for RPs, which are a measure of flux though the ribosome assembly pathway, most cancer cells display increased abundance of RIOK1, and RIOK1 abundance is more increased than the close relative RIOK2, or hCINAP, the human homolog for the ATPase Fap7, which both function directly prior to Rio1^3,62^ (Figure S5). While it remains unknown whether these changes in RIOK1 abundance play any role in promoting cancer progression by releasing miscleaved 18S rRNA-containing ribosomes into the translating pool, moving forward it will be important to understand how fluctuations in the relative concentrations of assembly factors affect QC during ribosome assembly and translation in human cells.

### Degradation of defective ribosomes

The 18S nonfunctional rRNA decay (NRD) pathway targets 18S rRNAs containing a mutation in its decoding center (18S:A1492C) for degradation. 18S NRD shares components with mRNA and protein QC (NGD and RQC, respectively), and all may be different outcomes of the same initiating event: ribosomes that stall on mRNA, leading to collisions with the tailing 80S ribosome to form an interface stabilized by Asc1^43,63^. Here we show that in the presence of mature ribosomes, functionally compromised miscleaved rRNAs are rapidly degraded in a pathway that shares many similarities with 18S NRD. Like NRD, it depends on translating ribosomes, Asc1, Dom34, Hel2, and Xrn1. This observation suggests that these collided disomes form the same interface as previously observed, which is then recognized by Hel2 to ubiquitinate Rps3^51,55^. Surprisingly, while important in the decay of non-functional 18S:A1492C rRNA, the decay of 3’-end miscleaved 18S rRNA is not dependent on Slh1, as Slh1 deletion affects neither cell growth nor the stability of miscleaved rRNA (Figure S2). This finding suggests that the Dom34/Hbs1-mediated splitting of ribosomes is the major way to disassemble nonfunctional 40S, while Slh1 is the major pathway for separating ribosomes collided due to imperfections in the mRNA^36,54,56^.

Furthermore, while Mag2 recognizes and ubiquitinates 18S rRNAs with a decoding site mutation (A1492C)^42^, it does not seem to be involved in the decay of miscleaved 18S rRNAs. We suspect that the differences arise from the specific complexes formed: while the A1492C mutation likely causes ribosomes to stall at the start site with an empty A-site, our data show that ribosomes with miscleaved 18S rRNAs translate. They might be slower moving or get stuck at specific codons, thereby causing collisions with correctly matured ribosomes within the ORF. Moreover, they are more likely to have a filled A-site, as the miscleaved 18S rRNA 3’-end is near the E-site, not the A-site. Previous work has shown that disassembly of collided ribosomes via the NGD pathway requires translation of at least 58 codons^51^, and that a different subset of factors targets collided disomes depending on whether the A-site is filled or not, and whether the leading ribosome is in the classic or hybrid state^36,54,64–66^. Thus, while the collision-dependent pathway we describe shares aspects of NGD and NRD, it also displays differences that seem to arise from the distinct molecular nature of the slow-moving collided disomes within the ORF. We suggest the name dysfunctional RNA decay (DRD).

### Implications for ribosome heterogeneity

Herein we show that ribosomes with altered function, even if they occur at low concentrations perturb the translation of correctly matured ribosomes, leading ultimately to the degradation of the ribosomes with altered function. In the case herein, the ribosomes with altered function contain miscleaved 18S rRNA, but the same should be true for ribosomes lacking modifications in the rRNA or RPs, or missing RPs that affect the ability of the ribosome to bind A- or P-site tRNA or to translocate efficiently, thereby slowing down elongation. The possibility of heterogeneous ribosomes arising from differences in rRNA modifications in functionally important regions has been raised and it has been speculated that they could affect global gene regulation^67–69^. The data here indicate that this is unlikely to be the case, because such heterogeneous populations, if they existed, would be rapidly homogenized via DRD, dysfunctional RNA decay. Importantly, ribosomes of altered function can persist if they sort themselves onto distinct classes of mRNAs, as we have shown for ribosomes containing and lacking Rps26^16^.

## Materials and Methods

### Yeast strains and cloning

*Saccharomyces cerevisiae* strains used in this study were obtained from Euroscarf, the Yeast Knockout Collection from GE Dharmacon (now Horizon Discovery Biosciences), or were made using PCR-based recombination^70^. Strain identity was confirmed by PCR, quantitative growth assays, and western blotting when antibodies were available. Site-directed mutagenesis was used to create mutations in plasmids, which were confirmed by sequencing. Plasmids were propagated in XL1 Blue competent cells. Yeast strains and plasmids used in this study are listed in Tables S1 and S2, respectively.

### rRNA 3’-RACE sequencing

#### Ribosome Purification

Gal::Pno1; Gal::Dim1 cells supplemented with plasmids encoding Dim1-E85A and either Pno1 or Pno1-KKKF were depleted of endogenous Pno1 and Dim1 by growth in YPD for 8 doublings at 25°C or 30°C. Cells were harvested between OD 1.0 – 2.0. Ribosomes were purified as previously described^71^. Ribosomal subunits were separated during purification and stored at -80°C as individual subunits. Due to concerns that reverse transcription through the Dim1-dimethylation site in 18S rRNA (m^6^_2_A1781 and m^6^_2_A1782) would pose complications, we used the dimethylation-incompetent Dim1-E85A mutation^44^ to allow us to sequence the final 40-60 nucleotides of 18S rRNA.

#### Library preparation and Illumina Sequencing

18S and 25S rRNA was isolated from 40S and 60S ribosomal subunits, respectively, by phenol-chloroform-isoamyl alcohol extraction. rRNA was treated with DNase I (New England Biolabs (NEB)) and size selected on a denaturing TBE-Urea-PAGE gel. rRNA was extracted from the gel by freeze-thawing the gel pieces in RNA extraction buffer (300 mM NaOAc, 1 mM EDTA, and 0.25% SDS) and ethanol precipitated. rRNA 3’-ends were dephosphorylated using rSAP (NEB) and ligated to either the Universal miRNA Cloning Linker (NEB; 25S rRNA samples and some 18S rRNA samples) or ligated to a pre-adenylated linker containing a UMI (unique molecular identifier, Integrated DNA Technologies (IDT), some 18S rRNA samples) using truncated T4 RNA ligase K227Q (NEB) to protect the 3’-end of the rRNAs from degradation and to identify the true rRNA 3’-end after sequencing (Table S3). Linkers were added in 2-fold excess of rRNA ends. 5’-ends of the DNA linkers and the rRNA were deadenylated by 5’-deadenylase and the excess linkers were degraded by DNA-specific RecJ_f_ exonuclease (NEB). rRNA was reverse transcribed using a linker-specific primer and Protoscript II RT (NEB) to generate the first strand of cDNA. After RNase H (NEB) treatment, the second strand was synthesized using an 18S or 25S rRNA-specific forward primer, a linker-specific reverse primer, and Q5 high fidelity DNA polymerase (NEB). The 18S rRNA-specific primer was designed to sequence either the last 41 or 62 nt of 18S rRNA, while the 25S rRNA-specific primer was designed to sequence the last 47 nt of 25S rRNA. The forward and reverse second strand synthesis primers contained partial P5 and P7 Illumina sequencing adapters, as indicated in Table S3. The cDNA was purified either on a denaturing TBE-Urea-PAGE gel and extracted by freeze-thawing in DNA extraction buffer (300 mM NaCl, 10 mM Tris-HCl (pH 8.0), and 1 mM EDTA) or cleaned up with AMPure XP beads (Beckman Coulter). The purified cDNA was amplified via PCR to generate the final libraries and add the complete P5 and P7 Illumina adapter sequences. The final size of the 25S cDNA libraries was about 194 bp, with a 64 bp insert, whereas the 18S cDNA libraries were either about 188 bp with 58 bp insert, or about 251 bp with 121 bp insert. Shorter 18S cDNA libraries were generated using the Universal miRNA Cloning Linker and the longer 18S cDNA libraries were generated using the UMI-containing linker. Library size and quality were assessed on an Agilent 2100 Bioanalyzer (Agilent Technologies). Validated libraries were pooled at equimolar ratios and loaded onto the NextSeq 500 flow cell. Primers for the reverse transcription and second strand synthesis reactions are listed in Table S3. All enzymes were bought from NEB and were used according to the manufacturer’s recommendations.

#### Bioinformatic processing

Demultiplexed and quality-filtered raw reads (fastq) generated from the NextSeq 500 were trimmed to remove Illumina adapter sequences with Trim Galore! (version 0.6.1) (Babraham Bioinformatics). Only reads containing the full linker sequence were retained, and the linker sequences were removed with CutAdapt (version 3.5)^72^ using Python version 3.8.3. Quality of trimmed reads was assessed using FastQC (version 0.11.4) (Babraham Bioinformatics). Trimmed reads were aligned to the *S. cerevisiae* 35S rDNA sequence (S288C) from the Saccharomyces Genome Database^73^ with Bowtie2 (version 2.2.9)^74^. SAMtools (version 1.1.0)^75,76^, BAMtools (version 2.4.0)^77^, and BEDtools (version 2.17.0)^78^ were used to identify reads aligning to the reference genome and to calculate the read depth or the fraction of reads cleaved at each nucleotide position. For 18S cDNA samples containing a UMI, the program UMI-tools (version 1.1.2)^79^ was used to extract the UMI sequence from each read prior to alignment. The extracted UMI sequence was then used to deduplicate reads after alignment but prior to calculating read depth and cleavage.

### Quantitative growth assay

NOY504 cells expressing plasmid-derived 18S rRNAs were grown overnight at 30°C in glucose dropout media, diluted for a day culture in the same media and grown for 3 hours at 30°C followed by 37°C for 5 hours until the cells reached mid-log phase. The cells were then inoculated into a 96 well plate in YPD media at OD 0.1. Cells expressing plasmid-derived 18S rRNAs with a BY4741 background (as indicated in Table S1) were grown overnight, diluted, and grown for an additional 3 hours to mid-log phase in glucose dropout media at 30°C. These cells were then inoculated into a 96 well plate in YPD media at OD 0.05. Cells expressing Pno1 or Pno1 mutants with and without Rio1 were prepared as previously described^7^. Cells were grown at 30°C or 37°C, as indicated in the figure legend, while shaking and the doubling times were measured in a Synergy 2 multimode microplate reader (BioTek).

### Sucrose density gradient analysis

Sucrose density gradient analysis and polysome profiling of whole-cell lysates followed by Western blotting and Northern blotting were performed as previously described^3,7^. All cells were grown to mid-log phase and then harvested. NOY504 cells expressing plasmid-derived 18S rRNAs were depleted of their endogenous ribosomes by growth at 37°C for 7 cell doublings prior to harvesting. Essential, endogenous proteins expressed under a Gal1 promoter were depleted in cells by growth in glucose dropout media for at least 12 hours prior to harvesting. The percent of 18S rRNA or 20S pre-rRNA in the polysome fractions was calculated by dividing the amount of 18S rRNA or 20S pre-rRNA in the polysome fractions (fractions 8-13) by the total amount of 18S rRNA or 20S pre-rRNA in all fractions (fractions 2-13). The percent of free Pno1 was calculated by dividing the amount of non-ribosome bound (free) Pno1 (fractions 1-2) by the total amount of Pno1 in all fractions (fractions 1-13).

### Northern Analysis

Northern blotting was carried out essentially as previously described (Strunk 2012), using probes listed in Table S3. For whole-cell RNA Northern blots, cells were grown in glucose dropout media at 30°C, except NOY504 cells which were grown at 37°C. 10 mL of cells at OD 0.5 were harvested, total RNA was extracted, and 5 µg of RNA per sample was used for the Northern blots.

### Antibodies

Primary antibodies against recombinant Rio1, Fap7, Rps10, Pno1, and Tsr1 were raised in rabbits by Josman or New England Peptide and tested against purified recombinant proteins and yeast lysates. The Rps8 antibody was a gift from G. Dieci and the Asc1 antibody was a gift from A. Link. The secondary antibody was anti-rabbit IgG conjugated to HRP from Rockland Immunochemicals. Blots were visualized using a BioRad ChemiDoc Imaging System.

### Protein expression and purification

Rio1 was purified as previously described^7^. Expression and purification of the maltose binding protein (MBP)-MS2 fusion protein was performed as described, with minor changes^80,81^. In brief, Rosetta DE3 competent cells transformed with a plasmid encoding His-MBP-tagged MS2 were grown to mid-log phase at 37°C in LB media supplemented with 2% glucose and the appropriate antibiotics. MBP-MS2 expression was induced by addition of 1 mM IPTG (isopropyl β-D-thiogalactoside), and cells were grown for another 5 hr at 30°C. Cells were lysed in 20 mM HEPES (pH 7.9), 200 mM KCl, 1 mM EDTA, 1 mM phenylmethylsulfonyl fluoride plus tablets of proteinase inhibitor mixture (Roche) by sonication on ice. The supernatant was applied to amylose resin and the MBP-MS2 protein was eluted in a buffer containing 20 mM HEPES (pH 7.9), 20 mM KCl, 1 mM EDTA, and 10 mM maltose. The protein was dialyzed into a buffer containing 20 mM HEPES (pH 7.9), 20 mM KCl, 1 mM EDTA, and 5 mM 2-Mercaptoethanol. The protein was then purified over a MonoQ column in a 20 mM - 1 M KCl gradient in 20 mM HEPES (pH 7.9), 1 mM EDTA, and 5 mM 2-Mercaptoethanol. Finally, the protein was dialyzed into a buffer containing 20 mM HEPES (pH 7.9), 100 mM KCl, and 10% glycerol. Protein concentration was determined with a Bradford assay.

### MS2-tagged RNA affinity purification (MS2-TRAP)

MBP-MS2-bound amylose resin was prepared in advance. First, 100 µL amylose resin was added to columns and washed 4 times with 1 mL H_2_O, and then 3 times with 1 mL MS2 storage buffer (20 mM HEPES (pH 7.4), 100 mM KCl, and 10% Glycerol). Purified MBP-MS2 protein (0.2 mg) was bound to amylose resin in 1 mL MS2 wash buffer (20 mM HEPES (pH 7.4), 200 mM KCl, 1 mM EDTA, and EDTA-free protease inhibitor (Roche)) by incubating at 4°C for 1 hr on a nutator. The amylose resin was then washed 4 times with 1 mL MS2 wash buffer and equilibrated with 2 mL ribosome lysis buffer (20 mL HEPES (pH 7.4), 200 mM KOAc, and 2.5 mM Mg(OAc)_2_). At this point, the MBP-MS2-bound amylose resin was ready for incubation with cell lysate.

Cells were grown in glucose dropout media at 37°C for 7 doublings and harvested at mid-log phase. Cells were then suspended in 0.5 mL/g of stringent ribosome lysis buffer (20 mL HEPES (pH 7.4), 200 mM KOAc, 2.5 mM Mg(OAc)_2_, 1 mM PMSF (phenylmethylsulfonyl fluoride), 1 mM DTT (dithiothreitol), 1 mM benzamidine, 1 µg/mL Leupeptin, 1 µg/mL Pepstatin, 10 µg/mL Aprotinin, and 1 mg/mL Heparin) and flash frozen in liquid nitrogen. Frozen cell pellets were additionally lysed by grinding into powder by mortar and pestle and thawed in 1 mL/g of stringent ribosome lysis buffer. Cell lysates were cleared and incubated with MBP-MS2-bound amylose resin in columns at 4°C for 1 hr on a nutator.

After ribosome binding, the resin was washed 4 times with 1 mL ribosome lysis buffer and equilibrated with 1 mL ribosome wash buffer (20 mM HEPES (pH 7.4), 100 mM KOAc, and 2.5 Mg(OAc)_2_). Finally, the MS2-bound ribosomes were eluted in two elution steps. The first elution was in 30 µL elution buffer (20 mM HEPES (pH 7.4), 100 mM KOAc, 2.5 Mg(OAc)_2_, and 15 mM maltose) after incubating the resin with elution buffer for 10 minutes. The second elution was in 200 µL elution buffer. For Western blot analysis, equal volume of 2x SDS-PAGE loading dye was mixed with the first elution and denatured at 95°C for 10 min before loaded onto an SDS-PAGE gel. Western blots were probed with the indicated antibodies.

### RNA binding assay

RNA binding assays were performed as previously described^19^. Briefly, ^32^P-ATP-labeled H44-D, H44- D+3, or H44- D-4 RNAs, named after the structural elements that mark their start and end points, were prepared by *in vitro* transcription in the presence of α-ATP. D+3 indicates that the RNA ends 3 nucleotides after the cleavage site D and D-4 indicates that the RNA ends 4 nucleotides before cleavage site D. The RNA transcription templates were PCR products containing the same promoter sequence at the 5’ end upstream of the H44 start site, and two 2’-O-methylated RNA nucleotides at the 3’-end to reduce T7 RNA polymerase’s non-templated nucleotide addition activity at the 3’-end of the RNAs, thus promoting uniformity at the 3’-ends of each RNA^61^. RNAs were then gel purified, eluted via electroelution, precipitated, and resuspended in water. RNAs were folded by heating for 20 min at 55°C in the presence of 50 mM HEPES (pH 7.5) and 10 mM MgCl_2_. Trace amounts of each radiolabeled RNA were incubated with varying concentrations of Rio1 with 1 mM AMPPNP in 50 mM HEPES (pH 7.5), 100 mM KCl, and 10 mM MgCl_2_ for 30 min at 30°C. Samples were loaded directly onto a running 6% acrylamide/THEM native gel to separate protein bound from unbound RNAs. After drying the gel, phosphorimager analysis was used to quantify the gel. Bound RNA was plotted against protein concentration and fit to Equation 1 to obtain apparent binding constants using GraphPad Prism version 8.4.3 (471) (GraphPad Software, La Jolla, California, United States, www.graphpad.com).

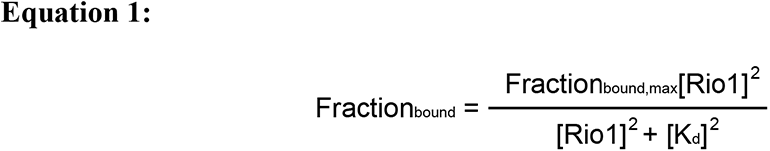

### Quantitative and statistical analysis

Quantification of Northern and Western blots was performed using ImageJ 1.53a (National Institutes of Health). Statistical analysis was performed using GraphPad Prism version 8.4.3 (471) (GraphPad Software, La Jolla, California, United States, www.graphpad.com). Statistical tests used and the number of samples (n) are indicated in the figure legends.

## Acknowledgements

This work was supported by National Institutes of Health (NIH) grants R01-GM086451 and R35-GM136323 and Howard Hughes Medical Institute (HHMI) Faculty Scholar grant 55108536 to K.K. We thank G. Dieci (Universita Degli Studi di Parma, Parma, Italy) and A. Link (Vanderbilt University, Nashville, TN) for the gifts of the anti-Rps8 and anti-Asc1 antibodies, respectively, and J. Doudna (University of California, Berkley, CA) for the gift of the plasmid encoding MBP-MS2. Purified MS2-MBP protein was a gift from M. Ferretti. We also thank M. Pipkin and M. Ferretti for helpful discussion on sequencing library preparation and assistance with bioinformatics, and H. Huang for providing processed RNA seq data from cancer samples.

## Author Contributions

Conceptualization: Melissa D. Parker, Katrin Karbstein

Funding acquisition: Katrin Karbstein

Performed experiments: Melissa D. Parker

Analyzed experiments: Melissa D. Parker and Adam J. Getzler

Project administration: Katrin Karbstein

Supervision: Katrin Karbstein

Writing: Melissa D. Parker, Katrin Karbstein

